# B-cell Follicles in the Secondary Lymphatic Tissues Act as a Critical Microanatomical Niche for Sustained Viral Replication, Virus-host Interaction and Damage During Chronic SIV Infection of Rhesus Macaques

**DOI:** 10.1101/2025.11.19.687931

**Authors:** Yilun Cheng, Miaoyun Zhao, Jackson Chen, Subhra Mandal, Mark G. Lewis, Qingsheng Li

## Abstract

Most people living with HIV-1 (HIV) are diagnosed during the chronic stage of infection, at which point viral setpoint has been established and a substantial immunopathologic damage in the secondary lymphatic tissues (LTs) has been induced. To better characterize virus-host interaction and immunopathologic damage in LTs, we investigated viral spatial distribution patterns and the corresponding LT structural alterations in SIV chronically infected rhesus macaques. Using RNAscope in-situ hybridization (RNAscope) and a combination of CODEX with RNAscope (Comb-CODEX-RNAscope), we demonstrated that in chronic infection, SIV viral RNA (vRNA) is predominantly resides within B-cell follicles with minimal presence in T-cell zones, and there is significant B-cell follicular damage, encompassing follicular hyperplasia, fibrosis, and disintegration of the germinal center. Our results reveal that the B-cell follicles act as a critical microanatomical niche for sustained viral replication, virus-host interaction and tissue damage during SIV chronic infection.

## Introduction

Most people living with HIV (PLWH) are diagnosed during chronic stage of infection. The progression of HIV infection from acute to chronic is a critical transition. While rapid viral replication and host-wide dissemination as well as initial CD4 T cell depletion, especially in gut, are characterized in the acute infection, which is usually defined as the first 3 months after infection (1, 2), the chronic phase features a persistent infection in the secondary lymphatic tissues (LTs) and a further decline in host immunity, including more CD4 T cell depletion and LT damage (3). LTs, such as lymph nodes and gut-associated lymphatic tissues (GALT), are important sites where immune cells reside, HIV replicates and persists, and virus-host interacts leading to CD4 T cell depletion, LT structure aberration, and immune dysfunction (4-7). Of note, GALT is the largest immunological organ in the body, of which ileum contains abundant lymphoid aggregates and Peyer’s patches that serve as key sites for viral replication and exhibit distinct immunologic microenvironments that influence the progress of chronic infection (8-11).

A better characterization of the HIV-host interaction in LTs during chronic infection can provide critical insights into the mechanisms of viral persistence and immunologic pathogenesis, and potential new therapeutic strategies (12). B-cell follicles in LTs function as a critical site for viral evasion of immunity due to a high abundance of follicular helper T cells (Tfh), which exhibit increased susceptibility to HIV infection, and, constitute a substantial target cells for persistent viral replication, even during antiretroviral therapy (ART) (13, 14). Simian immunodeficiency virus (SIV) infected rhesus macaques is the best available large animal model to study HIV pathogenesis, which enables comprehensive tissue sampling at specific time points that are unattainable in human studies (15). Here, we provide a comprehensive analysis of the viral distribution patterns and tissues damage in LN and ileal tissues during SIV chronic infection. We found that vRNA is primarily localized in the B-cell follicles, where there are B-cell follicular hyperplasia and fibrosis, indicating that B-cell follicles act as a critical microanatomical niche for sustained viral replication, virus-host interaction and damage during chronic SIV infection.

## Results

### SIV vRNA is mainly localized in B-cell follicles during chronic SIV infection

The secondary LTs are the major sites where HIV replication and persistence as well as virus-host interaction leading to immunopathologic damage and ultimately immunodeficiency without suppressive antiretroviral therapy (ART). LTs are anatomically divided into paracortex/ T-cell zones and B-cell follicles/B-cell zones, each has different immune cell composition and distinct immune functions. HIV has three viral forms, e.g., cell-free (V_CF_), cell-associated (V_CA_), and follicular dendritic cell (FDC)-trapped viruses (V_FDC_). To characterize three viral forms and their spatial distribution patterns during SIV chronic infection, we quantified vRNA and its cellular and compartmental localization in the mesenteric lymph nodes and ileal tissues from SIV chronically infected macaques using RNAscope in situ hybridization (RNAscope) and COMB-CODEX-RNAscope as we previously reported (16). During SIV chronic infection (180 dpi, n=5), the majority of vRNA was localized in B-cell follicles of LN tissues (92.47%) and lymphatic aggregates of ileal tissues (91.51%) (Figure 1) rather than in T-cell zones. In contrast, in early acute infection (10 dpi, n=3), the majority of viruses was localized in T-cell Zones of LN (54.59%) and lymphatic aggregates of ileal tissues (64.91%) (Figure 1). There was a substantial reduction of CD4 T cells in chronically infected LN and ileal tissues as comparison with acute infected macaques (Supplemental Figure1) as indicated by CD4/CD3 ratio. We quantified relative distributions in T-cell zones and B-cell follicles of total vRNA and vRNA^+^ CD4 T cells (V_CA_ in CD4 T cells) in SIV infected macaques using Comb-CODEX-RNAscope. We first separated vRNA signal channel into B-cell follicles versus T-cell zones using CD20 marker, then divided vRNA into several categories, e.g. V_CA_ in CD4 T cells or macrophages in T-cell zones or B-cell follicles as well as V_FDC_ in B-cell follicles by co-localizing vRNA signals with respective CD4 or CD68 markers (Figure 2B). In the B-cell follicles of chronically infected macaques, the majority of vRNA existed as V_FDC_, (vRNA signal manifested as diffused dots), nevertheless a small number of vRNA was colocalized with CD4 T cells, indicating ongoing Tfh cell infection in the B-cell follicles (Figure 2A, C & D). We compared how many percentages of total CD4 T cells were infected in the B-cell follicles versus T-cell zones as an indicator for CD4 T cell infectivity in the two sub-compartments (Figure 2 C). In addition, we quantified the frequency of virally infected CD4 T cells in given tissues area in the B-cell follicles versus T-cell zones, as an indicator for the relative density of infected CD4 T cells between B-cell follicles versus T-cell zones. We found there were significant higher of both CD4 T cell infectivity and the frequency of infected CD4 T cells in the B-cell follicles of ileal lymphatic aggregates and MES LN than in T-cell zones (Figure 2C-D). There is no significant difference in terms of CD4 cell infectivity and the frequency of infected CD4 T cells between LN tissues vs. ileal lymphatic aggregates (Figure 2C-D).

**Figure 1.**
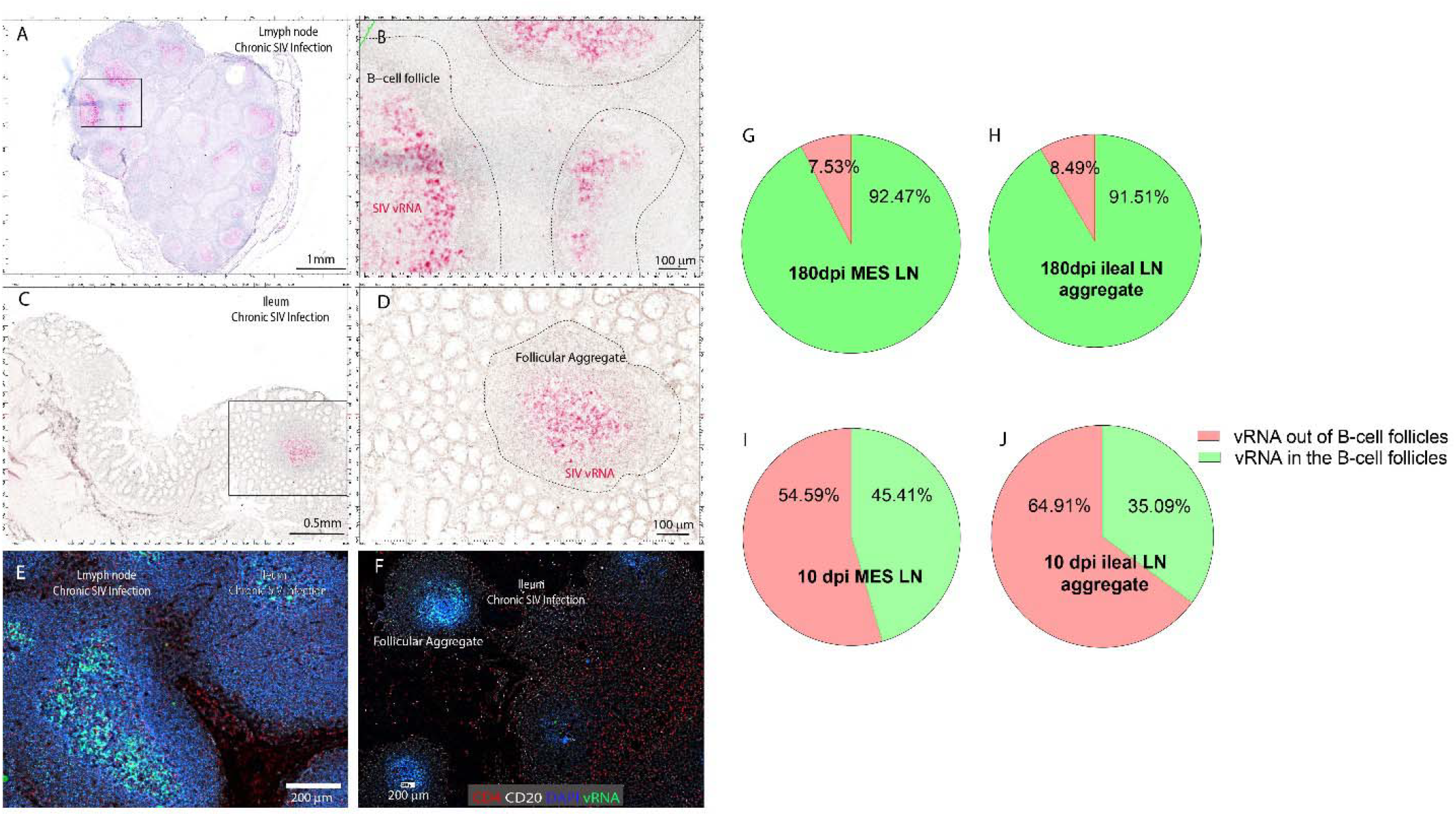
Viral RNA (vRNA) is mainly localized in the lymph node B-cell follicles and ileal follicular aggregates during SIV chronic infection. Representative micrographs showing SIV vRNA spatial distribution patterns in lymph nodes and ileum in SIV chronic infected macaques (A-F). SIV vRNA was detected using a combination of CODEX and RNAscope ISH (Comb-CODEX-RNAscope). **A-D** shows bright-field images of vRNA in red and nuclei in blue counterstained with hematoxylin, a higher magnification from highlighted boxed area in each figure was shown. **E-F** shows florescence channels of CD4 (red), CD20 (white), CD21 (Blue) and vRNA (green), **G-H** shows that vRNA was predominantly localized within the B-cell follicles of lymph node tissues (92.5%) and follicular aggregates of ileum (91.5%) in chronic SIV infection. **I-J** shows that in early infection the majority of vRNA was localized within the T-cell zones of lymph node tissues (54.59%) and follicular aggregates of ileum (64.91%).

**Figure 2.**
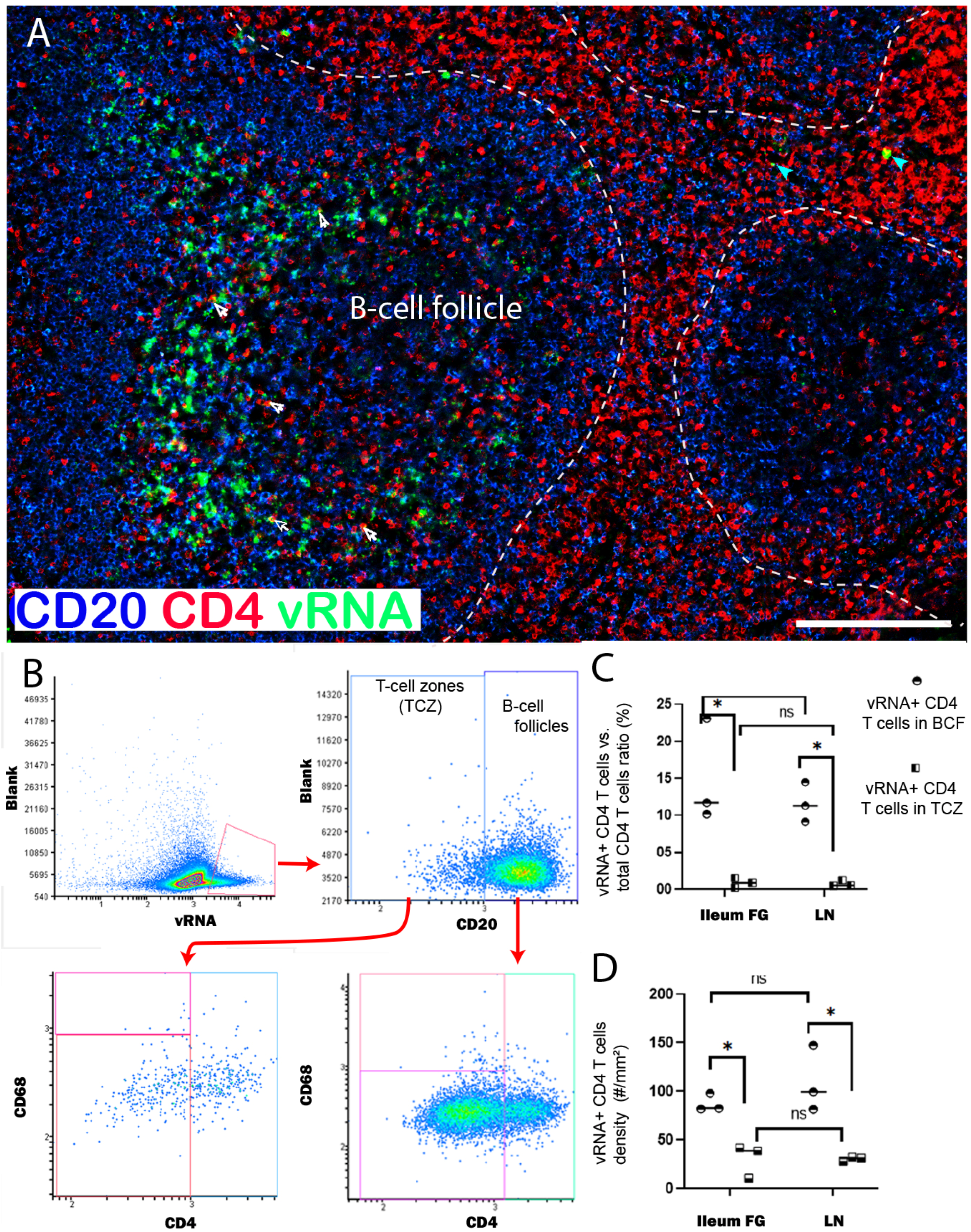
CD4 T cell infectivity and vRNA+ CD4 T cell density in B-cell follicles and T-cell zones. (A) Representative image of vRNA localization in lymph node tissue from a rhesus macaque chronically infected with SIVmac251 (BM41, 180 dpi), showing CD4 (red), CD20 (blue) and vRNA (green). White arrows indicate vRNA colocalized with CD4 T cells in the B-cell follicle, cyan arrows indicate vRNA colocalized with CD4 T cells in the T-cell zone. Scale bar indicates 200 um. (B) Gating strategy for vRNA in B-cell follicles versus T-cell zones, and colocalized cell types in each sub-compartment. (C) Scatter diagram shows CD4 T cell infectivity in B-cell follicles and T-cell zones. Each icon represents an individual resus macaque sample. (n=3, *P < 0.05, by T test) (D) Scatter diagram shows infected CD4 T cell density in B-cell follicles and T zones. Each icon represents an individual resus macaque sample. (n=3, *P < 0.05, by T test).

### B-cell follicular hyperplasia and fibrosis during chronic SIV infection

B-cell follicles are a key immune structure, where potent humoral immune responses take place through germinal center (GC) reactions. Moreover, follicular dendritic cells (FDCs) can capture complement-opsonized HIV virions through CD21/CD35 receptors, maintaining long-lived intact viruses within GCs (17). HIV Infection also induces IL-6/IL-21-mediated proliferation and accumulation of Tfh inside B-cell follicles (18). While providing substantial assistance to B cells, facilitating prolonged GC reaction and B cell proliferation (19), Tfh can be directly infected, such as through V_FDC_. We compared B-cell follicle size and fibrosis in SIV chronic infection vs. very early infection (10 dpi). We used B cell marker CD20 to demarcate, and measure expanded B-cell follicle size, proliferation marker Ki67 to evaluate GC activity and immune activation marker HLA-DR to evaluate immune activation in the B-cell follicles. We compared collagen levels in the B-cell follicles of LN tissues and ileal lymphatic aggregates in the chronic infection (180 dpi) versus early infection (10 dpi) and found there was exacerbated collagen deposition and fibrosis in the B-cell follicles of LN and ileal tissues in the chronically infected macaque tissues, but not in the acutely infected animals (Figure 3A), indicating increased collagen deposition and fibrosis and disrupted B-cell follicular architecture (Figure 3A & 4A-C). We also observed significant increase of Ki67 and HLA-DR protein levels in the B-cell follicles in chronically infected macaques as compared to acute infected animals (Figure 3B-C), indicating generalized immune activation and inflammation in the B-cell follicles during chronic infeciton. In addition, we observed that B-cell follicle size was drastically enlarged in chronic infected samples when compared to acute infected samples (Figure 4J). Also, in the SIV chronic infection, enlarged B-cell follicles also lose their intact structure, as enlarged/coalescence GCs or involuted GCs (Figure 4A-I).

**Figure 3.**
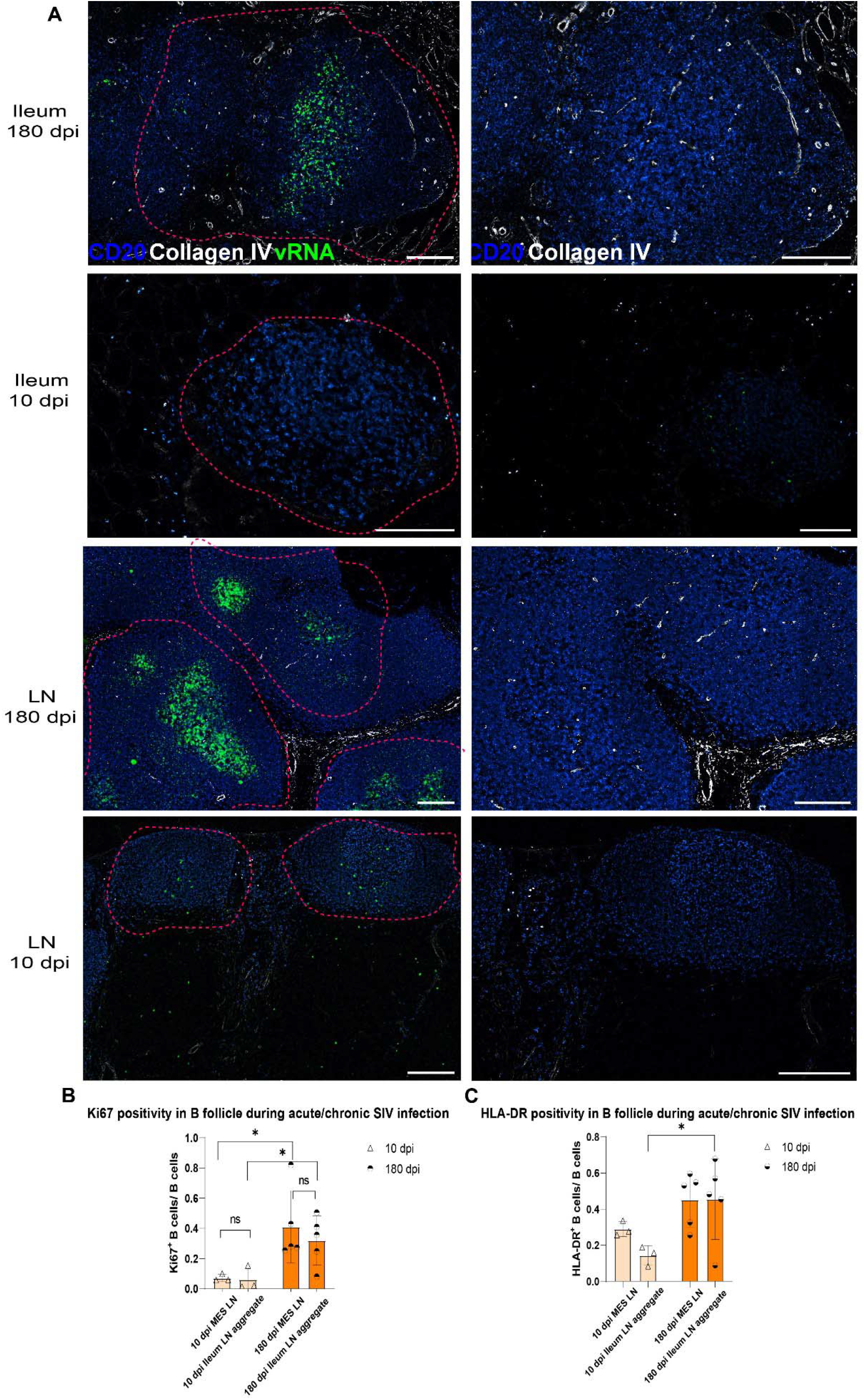
Collagen deposition, fibrosis, and immune activation in the B-cell follicles of LN tissues and ileal follicular aggregates during chronic SIV infection. (A) Representative images of collagen and vRNA locations in lymph node tissues or follicular aggregates of ileum from rhesus macaques infected with SIVmac251, showing Collagen IV (white), CD20 (blue) and vRNA (green). Scale bar indicates 200 um. (B) Scatter diagram shows Ki67 positive B cells in B-cell follicles. Each icon represents an individual resus macaque sample (n=3 acute SIV^+^, n=5 chronic SIV^+^, \**P* < 0.05, by *T* test). (C) Scatter diagram shows HLA-DR positive B cells in B-cell follicle. Each icon represents an individual resus macaque sample (n=3 acute SIV^+^, n=5 chronic SIV^+^, \**P* < 0.05, by *T* test).

**Figure 4.**
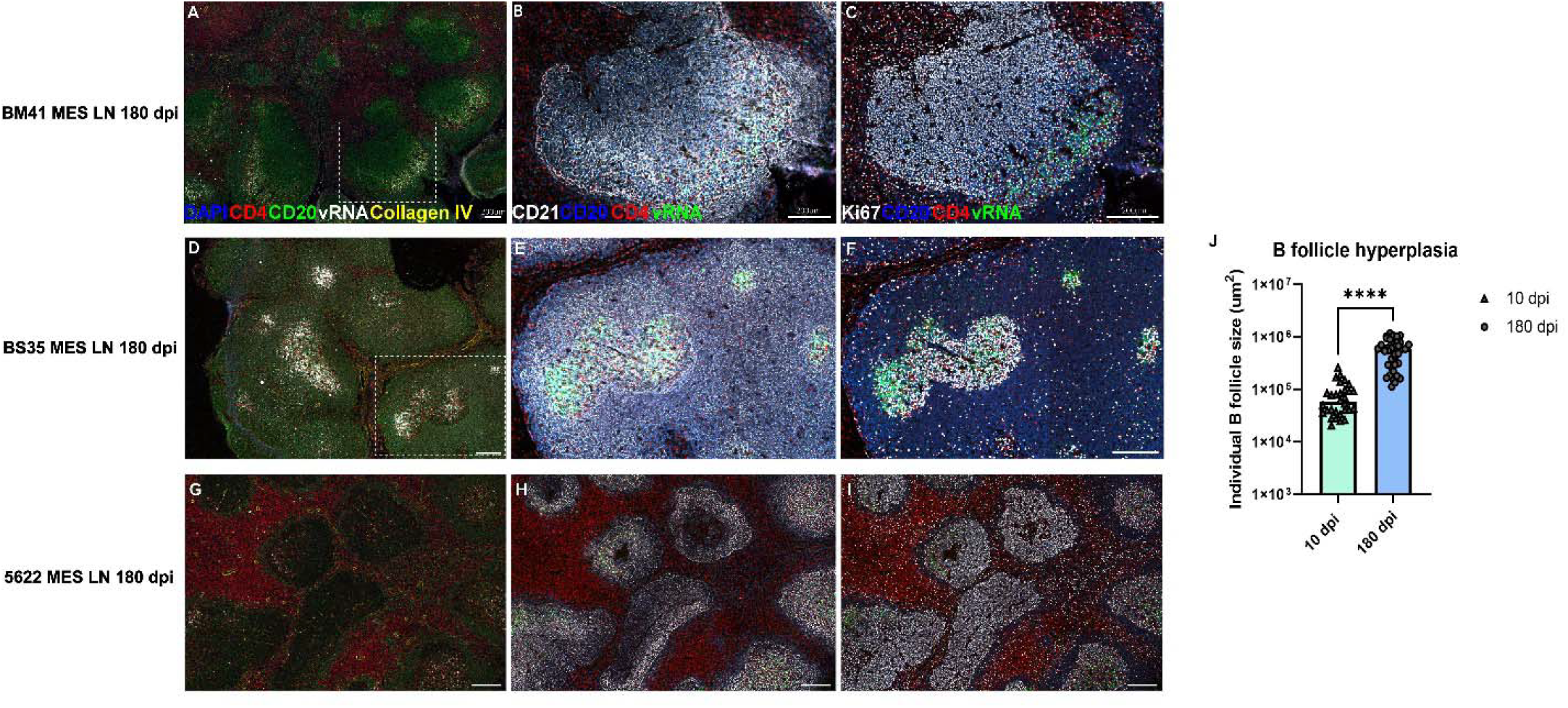
B-cell follicular hyperplasia during chronic SIV infection. Representative images of B-cell follicular hyperplasia in lymph node (LN) tissues from three rhesus macaques that were chronically infected with SIVmac251. Each row represents an individual macaque showing different channels of fluorescent markers. The scale bar equals 200um. **A-C** shows representative images of enlarged germinal center (GC), D-F shows a coalesced GC, and G-I shows several partially involuted GCs. (J) Scatter diagram shows averaged individual B-cell follicle size during chronic SIV infection as compared with early infection. Each icon represents an individual B-cell follicle from resus macaque samples (n=40 chronic SIV infection, n=30 acute SIV infection, ^+, +^, \**P* < 0.0001, by *T* test).

## Discussion

A comprehensive understanding of HIV-host interaction, viral existing forms, and structural alteration in B-cell follicles of LTs during HIV chronic infection is essential for the development, refining, and assessment of therapeutic strategies. In this study, we studied mesenteric lymph nodes and ileum during SIV chronic infections, because that they 1) contain enriched HIV targets cells (CCR5^+^ memory CD4^+^ T cells)(10, 20, 21), 2) are principal sites for viral replication, storage, and virus-host interaction (B-cell follicle, Tfh and V_FDC_)(11, 22), 3) undergo B-cell follicular structure changes that hinders immune reconstitution following antiretroviral therapy (ART) (23), and 4) facilitate gut microbial translocation that contributes to systemic immune activation (24). B-cell follicles are localized in the outer cortex of lymph node tissues and lymphatic aggregates of gut-associated lymphoid tissues (GALT). Primary B-cell follicles is predominantly consisted of B lymphocytes, which are differentiated upon antigen stimulation into the secondary B-cell follicles that are characterized by a B cell mantle encircling a core germinal center composed of centroblasts and follicular dendritic cells (FDC) (25). In this study, we used the Comb-CODEX-RNAscope technology to delineate the B-cell follicular, GC, and extrafollicular regions of lymph nodes and ileal lymphatic aggregates based on the spatial distributions of CD20 and Ki67 in follicular B cells and germinal centers (GCs), thereby enabled a relatively accurate evaluation of viral distribution patterns in B-cell follicles vs. T-cell zones and B-cell follicle damage during SIV chronic infection. We found that SIV vRNA loads are mainly concentrated within B-cell follicles during SIV chronic infection (180dpi) (Figure 1 A-C) rather than in T-cell zones. The higher loads of vRNA in B-cell follicles may be explained by several interconnected mechanisms 1) in the B follicle, T-follicular helper (Tfh) and regulatory (Tfr) cells are more susceptible and permissive to HIV/SIV infection than CD4 T cells in T-cell zones (18, 26-28), because of their higher expression of the CCR5 co-receptor (29), 2) a pro-survival state mediated by the anti-apoptotic protein BCL2 (26), and reduced expression of antiviral interferon-stimulated genes controlled by BCL6(18, 27), and 3) the B-cell follicle may serve as a sanctuary site for HIV/SIV infection due to ineffectiveness of CD8 T cells to clear infected cells. However, the final point remains controversial. Some researchers firmly believe that CD8 T cells cannot control viral replication in B-cell follicles since they are largely excluded from B-cell follicular sites (25, 30). Some studies indicated that the majority of HIV/SIV-specific cytotoxic CD8 T lymphocytes (CTLs) do not express the CXCR5 chemokine receptor, which is required for homing to B-cell follicles contributing to a much lower frequency of virus-killing cells inside the B-cell follicles compared to other areas of the LTs (26, 31, 32). Some studies indicated that even CD8 T cells managed to enter B-cell follicular regions, the low number and functional impairment (33, 34) lead to insufficient viral control, or the B-cell follicle microenvironment also promotes the development of regulatory CD8 T cell subsets and negatively contributing to viral clearance (33, 35). In contrast, other studies observed significant accumulation of CD8 T cells in B-cell follicles during chronic infection (36) and follicular CD8 T cells had cytolytic potential with high expression of granzyme B and perforin (36, 37). A recent study also identified T-follicular-like CD8 T cell responses (Tfc) in chronic HIV infection that were associated with both virus control and antibody isotype switching to IgG (38). In our study, we indirectly address this controversy by comparing CD4 T cell infectivity and infected CD4 T cell frequency in the B-cell follicles vs. T-cell zones (Figure 2 A-D). We observed a significantly higher rate of CD4 T cell infectivity and frequency in B-cell follicles than that in T-cell zones during chronic infection, indicating in B-cell follicles viral replication is less suppressed than T-cell zones.

We also observed that during SIV chronic infection there was B-cell follicular hyperplasia, neighboring follicles coalesced visually as GCs enlarge and FDC networks expand (Figure 4A). We also observed increased immune activation, collagen deposition and fibrosis in B-cell follicles (Figure 3). The collagen deposition and fibrosis can alter the stromal niche, impair normal cell trafficking and survival signals, and physically disrupt follicle architecture and function (39, 40). With chronic damage the FDC network that scaffolds GCs becomes fragmented or depleted; without an intact FDC network GCs cannot maintain affinity maturation or normal B cell survival, accelerating GC collapse and follicles involute and may be replaced by fibrotic scar tissue (Figure 4A-I). The surviving follicular structures are also frequently abnormal: architectural disruption, persistent trapped viruses on FDCs, and fibrosis (Figure 3A and 4A-I).

In conclusion, our study showed that B-cell follicles serve as a critical site for SIV replication and persistence, which contributes to structural and functional damage of B-cell follicles, resulting in compromised humoral immunity and systemic deficiencies in anti-SIV immunity.

## MATERIALS AND METHODS

### Rhesus Macaque Tissues

Archival fixed lymphatic tissues from SIVmac251-infected adult rhesus macaques from our previously published research were used(41, 42). Rhesus macaques were inoculated intra-rectally with SIVmac251 (3.1 x 10^4 TCID50) and subsequently euthanized at various time points post-inoculation, while uninfected controls were maintained without virus inoculation. Lymph node tissues were fixed using SafeFix II (Cat# 23-042600, Fisher Scientific), 4% paraformaldehyde, or neutral-buffered formalin and subsequently embedded in paraffin.

### Antibody-oligonucleotide conjugation

To identify a rhesus macaque reactive CD3 antibody for CODEX, an anti-human CD3 antibody in a carrier-free PBS solution (Clone#: SP162, Cat#: ab245731, Abcam) was conjugated with the barcode-oligonucleotide (Cat#: 5350002, Akoya) utilizing the CODEX conjugation kit (Cat#: 7000009, Akoya) in accordance with the CODEX conjugation manual.

### RNAscope ISH

RNAscope in situ hybridization (ISH) was conducted in accordance with the manufacturer’s procedure and our previously reported methodology(43, 44) with minor adjustments. All processes, including deparaffinization, hydration, and antigen retrieval, were performed to reduce RNase contamination. Tissue slides were deparaffinized by successive heating on a hotplate at 60°C for one hour, followed by immersion in xylene for five minutes twice, and subsequently rehydrated with progressively reduced concentrations of ethanol and DEPC water. Followed by incubation in 3% hydrogen peroxide for 10 minutes at room temperature. Following washing with mili-water (AQUA SOLUTION, 2700PRD), the tissue coverslips underwent antigen retrieval by boiling for 15 minutes in RNAscope® Target Retrieval Reagent solution (Cat # 322000, ACD) within a 50ml glass beaker on a hotplate (Thermo Fisher, SP88857104) to reverse cross-linking. Tissue coverslips were subjected to treatment with RNAscope® Protease Plus (Cat # 322330, ACD) at 40°C for 20 minutes, followed by hybridization with RNAscope® Probe-SIVmac239 (anti-sense, Cat# 312811, ACD) at 40°C for 2 hours. The signals were amplified and detected using the RNAscope® 2.5 HD assay-Red kit (Cat# 322360, ACD), allowing visualization of fast red in both standard light and far-red fluorescence channels. The RNAscope® negative control probe-DapB (Cat# 310043, ACD) served as the negative control.

### Combination of CODEX immunostaining with RNAscope ISH

The combination staining of CODEX-ISH was conducted in accordance with the CODEX user handbook(45) and our previous established methodologies(16). Six-micrometer tissue sections were prepared using a Leica RM2235 rotary microtome and subsequently mounted onto poly-lysine coated coverslips. In the CODEX and subsequent RNAscope approach, pretreatment involving 3% hydrogen peroxide incubation, antigen retrieval was conducted prior to the CODEX cycle as previous mentioned in RNAscope staning part, and the proteinase digestion step was following the final CODEX cycle. Following antigen retrieval, the tissue-coverslips were incubated with a mixture of DNA-barcoded antibodies for 3 hours at room temperature. Following three washes in PBS for 2 minutes each to eliminate unbound antibodies, the tissue-coverslips underwent post-fixation with 1.6% PFA for 10 minutes at room temperature, were subsequently washed, and then post-fixed with ice-cold methanal for 5 minutes before another wash. The tissue coverslips can be stored in the storage buffer for five days at 4°C or immediately loaded into the CODEX instrument for multiple cycle immunostaining. The CODEX fluidics and Keyence microscope were configured utilizing CODEX Instrument Manager (CIM) software and Keyence software in accordance with the manufacturer’s protocol. The master mix of fluorescently labeled oligonucleotide probes, which corresponded to DNA-barcoded antibodies for multiple cycle immunostaining, was prepared in a 96-well plate in accordance with the CODEX manual. The CODEX immunostaining process comprised three steps: the addition of nuclear staining DAPI, application of Atto550-, Cy5-, and Cy7-fluorescent reporters, followed by imaging and subsequent removal of the fluorescent reporters. Two blank cycles, consisting solely of DAPI nuclear stain and devoid of any fluorescent reporters, were conducted to assess autofluorescence levels and to subtract background using CODEX processor software. After removing the tissue-coverslip from the CODEX instrument, the procedure can proceed with the single RNAscope protocol starting from the proteinase digestion step. Following the RNAscope procedure, the tissue-coverslip was stained with DAPI (1:2000) for 5 minutes at room temperature, washed, and subsequently loaded into the CODEX instrument for the acquisition of RNAscope fluorescent images. To integrate RNAscope images with CODEX images, the Keyence microscope was configured using the Keyence software as previously described, and a new blank cycle was introduced, with the Cy5 channel designated for RNAscope. Following the acquisition of the RNAscope fluorescent image, the image data was transferred to the designated CODEX file location, where the raw data from CODEX and RNAscope underwent processing using CODEX processor software.

## ACKNOWLEDGMENTS

This work was supported by the National Institutes of Health, R01 DK087625.

---

**Supplemental Figure 1.**
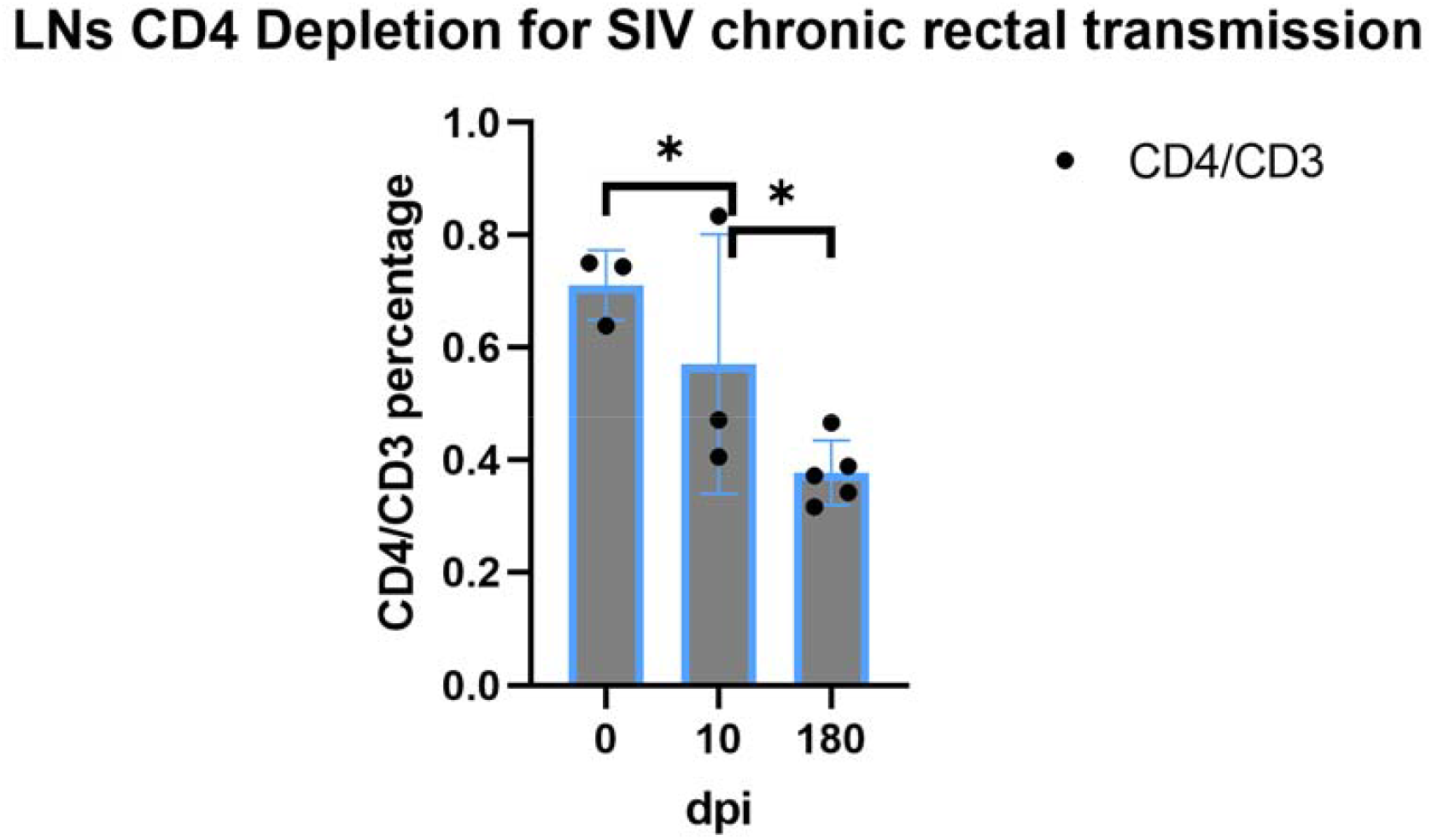
CD4 T cell depletion in chronic SIV rectal transmission. Scatter diagram pooled data showing CD4 T cell depletion during SIV rectal transmission. Each icon represents an individual resus macaque sample (n=3 non-infected control and acute SIV^+^, n=5 chronic SIV^+^, * *P* < 0.05, by *T* test).

## Notes

### Competing Interest Statement

The authors have declared no competing interest.

